# The Rho effector ARHGAP18 coordinates a Hippo pathway feedback loop through YAP and Merlin to regulate the cytoskeleton and epithelial cell polarity

**DOI:** 10.1101/2024.11.26.625473

**Authors:** Emma C. Murray, Gillian M. Hodge, Khanh Pham, Leighton S. Lee, Cameron A.R. Mitchell, Yongho Bae, Andrew T. Lombardo

## Abstract

The organization of the cell’s cytoskeletal filaments is coordinated through a complex network of signaling cascades activated by both internal and external cues. Two major actin regulatory pathways are signal transduction through Rho family GTPases and growth and proliferation signaling through the Hippo pathway. These two pathways define the actin cytoskeleton, controlling foundational cellular attributes such as morphology and polarity and are hijacked to promote proliferation and motility in aggressive cancers. In this study, we use human epithelial cells to investigate the interplay between the Hippo and Rho Family signaling pathways, which have predominantly been characterized as independent actin regulatory mechanisms. We identify that the RhoA effector, ARHGAP18, forms a complex with the Hippo pathway transcription factor YAP to address a long-standing enigma in the field. Using super resolution STORM microscopy, we characterize single-filament-level changes in the actin cytoskeleton that arise from CRISPR/CAS9 knockout of ARHGAP18. We report that the loss of ARHGAP18 results in cytoskeletal alterations driven by both dysregulated RhoA signaling and aberrant nuclear localization of YAP. These findings indicate that the Hippo and Rho family GTPase signaling cascades are temporally and spatially coordinated in their regulation of the actin cytoskeleton.

## Introduction

Extracellular signals are transduced through the cell membrane, in part through receptor ligand activation of Rho family GTPases. Rho family proteins bind and cycle between an active, guanosine triphosphate (GTP) bound, and an inactive guanosine diphosphate (GDP) bound state. This cycling has traditionally been referred to as a ‘molecular switch’ that turns on or off major actin regulatory components (Hall and Nobes, 2000). These components include the actin-based molecular motor non-muscle myosin 2 (NM-2) through Rho Associated Kinase (ROCK) or the actin severing protein cofilin, through LIM kinase (Julian and Olson, 2014). Recent understanding has advanced the ‘on/off’ characterization of Rho GTPases to a more nuanced regulatory environment where both GTP and GDP bound states promote molecular interactions temporally and spatially (Denk-Lobnig and Martin, 2019). None-the-less, Rho family GTPases are well established as essential cellular signals for actin nucleation, polymerization, and organization into higher-order branched or bundled structures (Hall, 1998). These signals serve to order the cell into polarized domains, defining the shape of the cell, the morphological structures present (e.g. microvilli, filopodia, etc.), and the cell-cell contacts required to build tissues.

Recently, a conserved mechanism of actin regulation has emerged where domain-specific regulation by Rho effectors is controlled on a minute temporal and spatial scale (Jackson et al., 2024; Landino et al., 2021; Sepaniac et al., 2023; Swider et al., 2022). Our group recently characterized an epithelial specific RhoA signaling pathway involving ARHGHAP18 organizing at the apical surface of polarized cells on the scale of 100nm (Lombardo et al., 2024). Separately, we showed in human colorectal CaCo-2 cells that the RAB GAP Epi64A localizes to an approximately 75nm region at the base of microvilli to regulate the terminal web of actin controlling cell shape (Miller et al., 2022). Collectively, these findings indicate that from flies to humans, the cell regulates Rho family GTPases on the scale of each individual actin structure to build the correct structure at the correct time. In the progress of our studies, we discovered an unexpected phenotype in cells genetically lacking ARHGAP18. These cells exhibited a basal membrane actin phenotype opposite to what would be expected from the loss of a RhoA GAP, challenging the emerging model in the field.

ARHGAP18 shows a strong specificity for accelerating the hydrolysis of GTP of RhoA in epithelial cells (Maeda et al., 2011) but also exhibits increased activity toward RhoC in endothelial cells (Chang et al., 2014). RhoA-GTP regulates the actin cytoskeleton by promoting multiple downstream kinase pathways. Of these, two kinases are required for the formation actin organization in polarized epithelial cells: Rho-associated protein kinase (ROCK) and the homologues Lymphocyte Oriented Kinase/ STE20-like Kinase (LOK/SLK). These kinases increase actin bundling in both stress fibers through non-muscle myosin-2 (NM-2) and microvilli through activation of the Ezrin, Radixin, Moesin (ERM) family in concert with recruitment of additional microvilli specific components (Gaeta et al., 2021; Morales et al., 2023; Zaman et al., 2021). RhoA GAPs limit these effects by accelerating the transition of RhoA, negatively regulating them toward its inactivated GDP-bound state. Thus, increased RhoA GAP activity canonically scales inversely with the formation of bundled actin filaments. However, ARHGAP18 knockdown results in the disorganization of stress fibers in human cells (Lay et al., 2019). The fly homolog of ARHGAP18 was similarly reported to positively regulate actin bundling through Rho1 and was subsequently named conundrum (Conu) due to this unexpected phenotype (Neisch et al., 2013). Astonishingly, knockdown of ARHGAP18 in human 3D spheroids was reported to reduce NM-2 activation and resulted in a loss of stress fibers (Porazinski et al., 2015).

To explain these results, ARHGAP18 is hypothesized to act as a non-traditional RhoA GAP where genetic screening linked its enigmatic function to the cytoskeletal transcription factor Yes-associated protein (YAP) (Porazinski et al., 2015). Additionally, in ARHGAP18 deleted mice, YAP inappropriately localizes to the nucleus (Coleman et al., 2020). YAP and the transcriptional co-activator with PDZ-binding motif (TAZ) are the downstream effectors of the Hippo pathway. Hippo signaling originates with mechanosensation and nutrient availability at apical and junctional membranes in part through the ERM-related protein Merlin (also known as neurofibromin 2 or NF2) (Harvey et al., 2013; Reginensi et al., 2016). Hippo signaling shuttles YAP and TAZ between the cytoplasm and nucleus. When localized to the nucleus, YAP/TAZ promotes the activation of cytoskeletal transcription factors associated with cell proliferation and actin polymerization. Phosphorylation of YAP at the S127 residue targets it to the cytoplasm for eventual degradation. Alternatively, the activation of YAP and TAZ is pervasive in human malignancies (Zanconato et al., 2016). The combined results of the studies linking ARHGAP18 to YAP/TAZ reported both upstream and downstream regulation, indicating a possible feedback loop. However, the mechanism of this potential relationship remained unknown.

In this manuscript, we characterize the aberrant actin networks of Jeg-3 human epithelial cells genetically lacking ARHGAP18. Using super resolution STORM to resolve the actin filament network to a resolution of <40nm, we report that loss of ARHGAP18 results in a near total loss of basal actin bundles, including stress fibers and filopodia. We characterize the alterations to the kinases downstream of RhoA signaling regulating basal actin organization. Our results indicate that the defective basal actin organization results from both RhoA regulation through canonical GAP activity and separately through ARHGAP18’s regulation of YAP localization. We report that ARHGAP18 forms a stable complex with YAP, which regulates its cytoplasmic to nuclear shuttling to control basal actin organization. Further, ARHGAP18 binds to Merlin, enabling ARHGAP18 to act as a feedback system for the Hippo signaling pathway. This mechanism allows the cell to coordinate the Hippo and Rho family signaling pathways to regulate the actin cytoskeletal organization in response to both internal and external cues.

## Results

### ARHGAP18 Regulates Non-Muscle Myosin-2 Through a RhoA Independent Mechanism

Our group recently characterized the regulation of microvilli through direct binding of ARHGAP18 to Ezrin, where we produced and validated CRISPR/CAS9 knock out of ARHGAP18 in human epithelial Jeg-3 cells (Fig. 1A) (Lombardo et al., 2024). In microvilli, ARHGAP18 acts to suppress active RhoA leading to reduced ROCK activation and lower MLC phosphorylation. Surprisingly, cells lacking ARHGAP18 and cells overexpressing ARHGAP18 both show increased phosphorylation of myosin light chain (Fig. 1B). Total expression levels of NM-2, total MLC, and actin are unchanged across these conditions (Fig. 1B). We hypothesized that the overexpression of ARHGAP18 was regulating the downstream RhoA kinase ROCK independent of its GAP activity. To probe this possibility, we tested if the alternative signaling cascades downstream of ROCK were functioning properly. Using phosphospecific antibodies for western blotting, we detected that phosphorylation of LIM kinase (pLIMK) and cofilin (pCofilin) exhibited only minor changes in the absence of ARHGAP18 (Fig. 1C, D). While the abundance of phosphorylated LIMK appeared slightly increased in the KO cells, it did not rise to the level of statistical significance. Unlike pMLC, over-expression of ARHGAP18 resulted in only a modest decrease in phosphorylation of LIM kinase and Cofilin (Fig. 1D), which was not significantly different than WT cells. We concluded that in ARHGAP18 altered cells LIMK, Cofilin, and MLC were in part regulated through the RhoA GAP activity of ARHGAP18. However, MLC in the ARHGAP18 overexpression was regulated through a GAP-independent mechanism.

**Figure 1:**
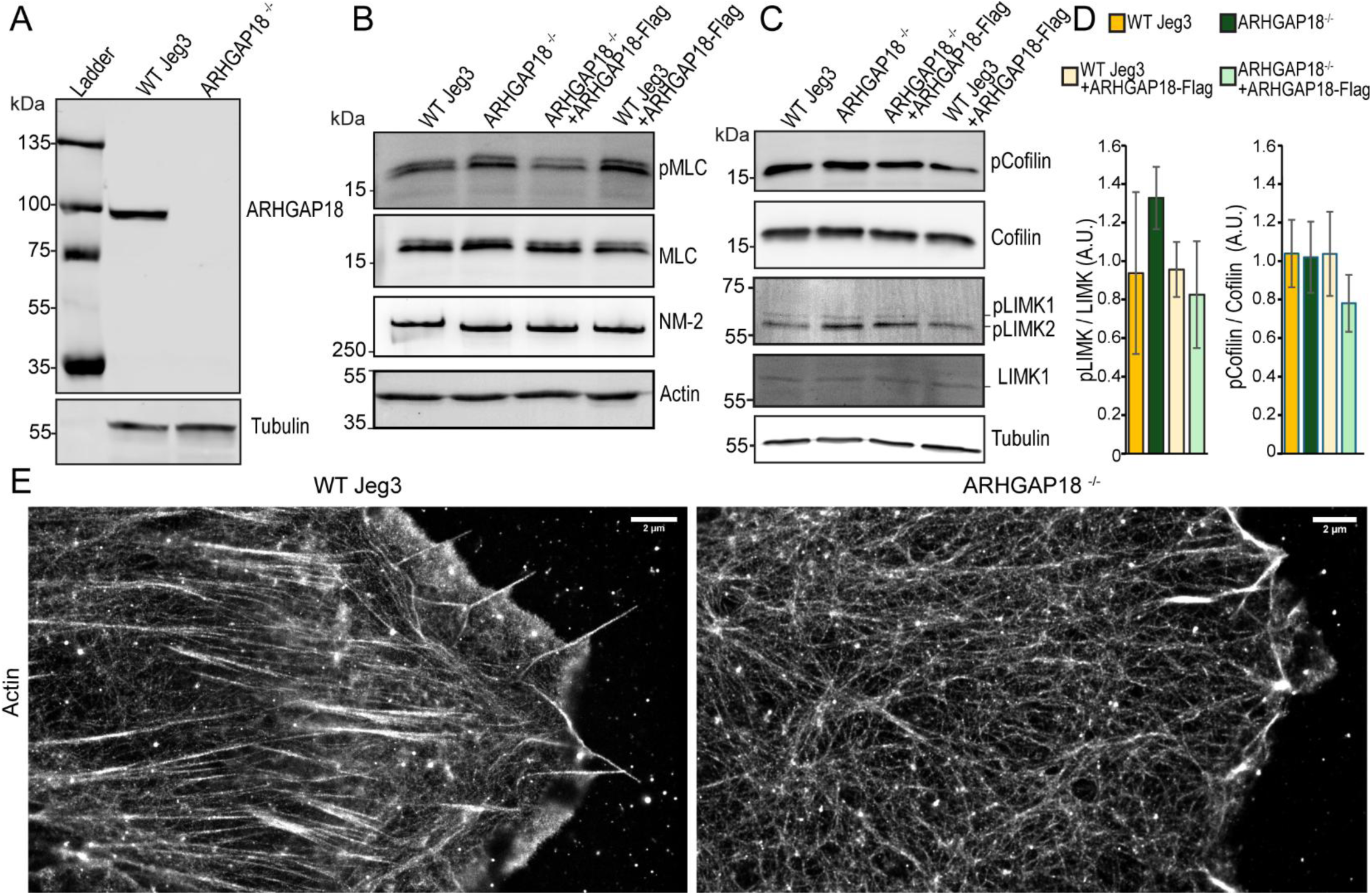
Characterization of ARHGAP18 Knockout in Human Epithelial Cells. A) Western blot of ARHGAP18 and Tubulin in WT Jeg-3 cells and ARHGAP18^-/-^ knockout (KO) cells. B) Western blot of Non-muscle Myosin 2 (NM-2), Myosin Light Chain (MLC), and phosphorylated MLC (pMLC) in WT and KO cells with tagged ARHGAP18 rescue or overexpression. C) Western blot of total cofilin and LIM kinase vs. phosphorylated fractions of each in the same conditions as (B). D) Quantification of western blot intensity of the ratio of phosphorylated fraction over total for cofilin and LIMK in the conditions from (C). Bars indicate Mean±SEM; All conditions non-significant by t-test (p>0.05) n=3 E) STORM reconstructions of phalloidin stained Actin in fixed Jeg-3 WT and Jeg-3 ARHGAP^-/-^ cells. Scale bar 2µm.

### Cells Lacking ARHGAP18 have disorganized basal actin and a loss of focal adhesions

Increased pMLC in the absence of ARHGAP18 is expected to result in increased stress fiber formation through the activation of NM-2. However, multiple independent investigations, including our own, had identified the loss of stress fibers in cells with reduced or lacking ARHGAP18 (Lay et al., 2019; Lombardo et al., 2024; Neisch et al., 2013; Porazinski et al., 2015). However, the true structure of actin filaments is well below the diffraction limited resolution of light, which forms dense arrays of filaments at the plasma membrane cortex interface. To characterize the actin filament organization at the basal surface of cells lacking ARHGAP18, we employed Stochastic Optical Reconstruction Microscopy (STORM) to resolve the individual actin architecture to a spatial X-Y resolution of less than 40nm. WT Jeg-3 cells showed numerous stress fiber bundles at the basal surface of the cell (Fig. 1E). Actin filaments transitioned to lateral filaments running parallel to the plasma membrane and eventually to a canonical leading edge comprised of both cortical branched individual filaments and filopodia at the cell’s plasma membrane. The orientation of the stress fibers and filopodia generally aligned in the axis going from cell center to cell exterior. In contrast to the WT actin organization, cells lacking ARHGAP18 exhibit a striking absence of both stress fibers and filopodia (Fig. 1E). The actin of ARHGAP18 KO cells was organized into small bundles and long individual filaments. Notably, the orientation of the ARHGAP18 KO cells’ actin was substantially less aligned in the general axis going from cell center to cell exterior seen in the WT cells.

To further characterize the basal actin phenotype in cells devoid of ARHGAP18 further, we used immunofluorescent staining of NM-2B using spinning disk confocal. Stress fibers from WT cells showed actin colocalized with repeating striations of NM-2B indicative of active, contractile fibers (Fig. 2A). NM-2B still colocalized to actin in ARHGAP18 deficient cells but organized into smaller more diffuse puncta. We previously reported that ARHGAP18-deficient cells were nearly twice as stiff as WT cells when measured by force indentation using atomic force microscopy (Lombardo et al 2024). Given these data combined with our western blotting of pMLC, we concluded that NM-2B was still active in cells lacking ARHGAP18 and capable of forming into contractile bundles but that the force-producing activity was not organizing the actin into large stress fiber bundles seen in the WT cells.

**Figure 2:**
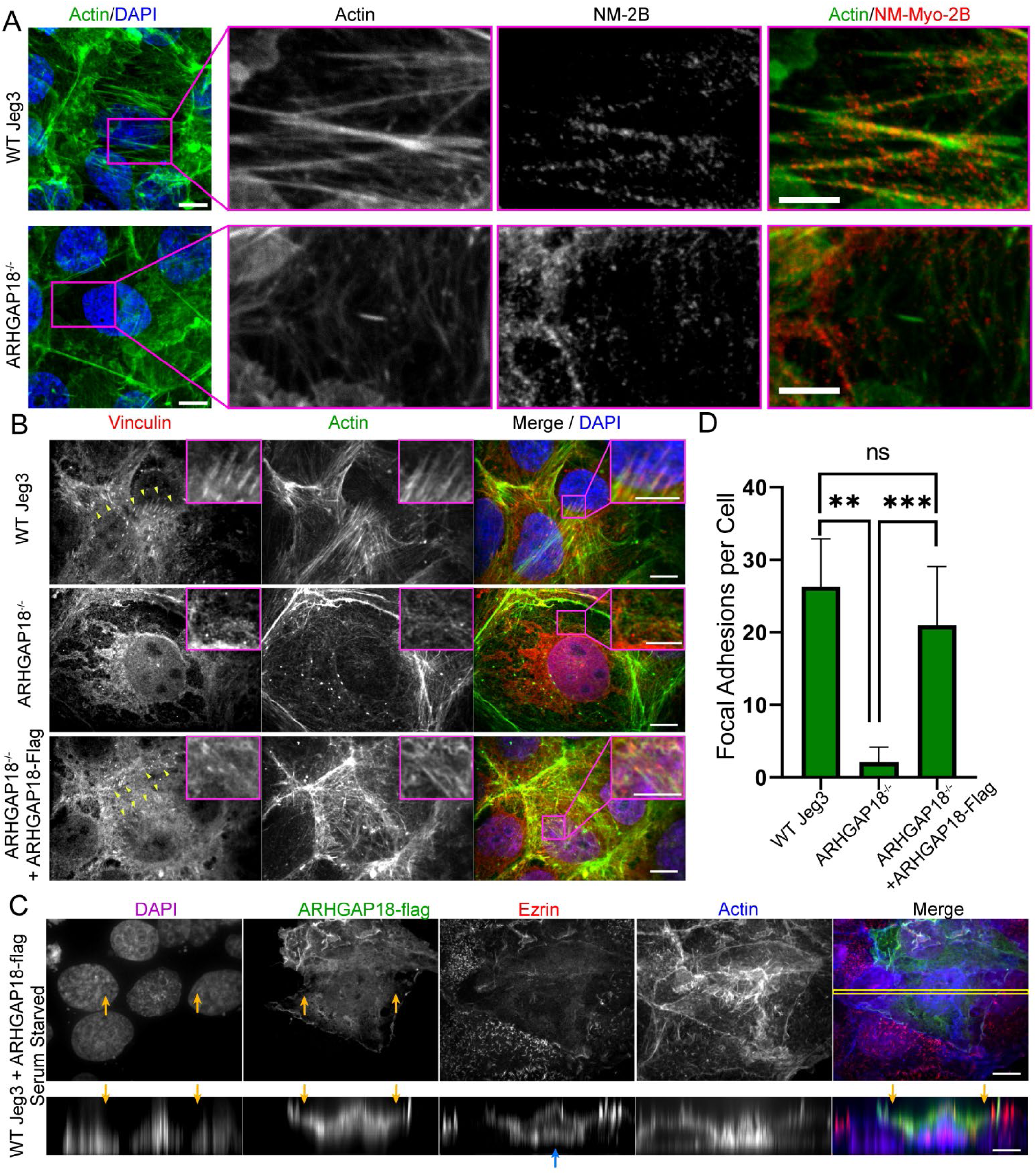
ARHGAP18 Binds Merlin and Effects NM-2 and Vinculin at the Basal Surface. A) Immunofluorescent imaging of actin and Non-Muscle Myosin-2B (NM-2B) in WT and ARHGAP18^-/-^ cells. Scale bar 10µm; inset 5µm B) Immunofluorescent imaging of vinculin and actin in WT and ARHGAP18^-/-^ cells and ARHGAP18-flag rescue cells. Focal adhesions co-localizing actin and vinculin highlighted with yellow arrowheads. Scale bar 10µm. Inset scale bar 5µm. C) Spinning disk confocal immunofluorescent imaging of WT Jeg-3 cells overexpressing ARHGAP18-flag. Actin to plasma membrane crosslinker ezrin localizes exclusively to the apical plasma membrane. Lower image row shows the X-Z dimension of the same image showing the invasion of the overexpressing cell onto neighboring cells. Yellow arrows indicate the sections of the overexpressing cell invading over neighboring cells’ nucleus. Blue arrow indicates two stacked apical membranes by ezrin staining with the lower membrane from WT cells underneath the upper membrane from an ARHGAP18 overexpressing cell. Scale bars 10µm. D) Quantification of the number of vinculin-stained focal adhesions per cell. Bars are Mean ± SEM. conditions are significant by Mann-Whitney test; WT n=average of 4 replicates, ARHGAP18^-/-^ n=average of 7 replicates, ARHGAP18^-/-^ +ARHGAP18-Flag n= average of 7 replicates (**= p ≤ 0.01, *** = p ≤ 0.001, n.s.= not significant).

To investigate why basal actin organization was disrupted despite the activation of NM-2 in ARHGAP18 KO cells, we investigated the presence of actin to basal-membrane contact proteins. Immunofluorescent imaging of Vinculin in WT cells indicated that stress fibers were tethered to the plasma membrane at the basal surface (Fig.2B). Imaging of ARHGAP18 ^-/-^ cells showed mislocalization of vinculin with no focal adhesions observed. Expression of full-length human ARHGAP18-flag in ARHGAP18 ^-/-^ cells rescued both actin basal actin bundles and vinculin focal adhesion localization (Fig 2B). We sought to characterize the defects of pMLC overactivation and vinculin mislocalization in cells overexpressing ARHGAP18-flag. However, we discovered that WT cells overexpressing ARHGAP18 would often break from the surrounding Jeg-3 cell’s monolayer to form three-dimensional stacks of cells (Fig. 2C). This characteristic was not typically seen in monolayered WT and ARHGAP18^-/-^ cells (Fig. 2A). To investigate this invasive phenotype, we used transient transfection of ARHGAP18-flag into WT cells which allowed us to compare the behavior of WT cells side-by-side with ARHGAP18 overexpressing cells. Spinning disk confocal Z-dimensional slicing of the actin on these cells indicated that large basal actin bundles and stress fibers were present in the ARHGAP18 overexpression cells (Fig. 2C). We observed that cells overexpressing ARHGAP18 invade their neighboring cells monolayer climbing almost entirely on top of the adjacent WT cells (Fig. 2C yellow arrows). These data support the conclusion that ARHGAP18 acts to regulate basal and junctional actin through Rho-independent mechanisms.

### Cells Lacking ARHGAP18 Have Transcriptional Changes in Actin and Proliferation Regulators

We hypothesized that the invasive and cytoskeletal phenotypes observed at the basal surface of cells devoid of ARHGAP18 may be a result of changes in regulation at the transcriptional level either directly through RhoA signaling or through an additional mechanism specific to ARHGAP18. To test this hypothesis, we performed bulk RNA sequencing (RNAseq) on WT-Jeg-3 and ARHGAP18 KO cells. We confirmed expressional changes between WT and ARHGAP18 KO cells using PCA analysis (Fig 3A). The analysis revealed two distinct clusters corresponding to WT Jeg-3 and ARHGAP18 KO samples, suggesting clear transcriptomic differences between the two groups. A total of 889 differentially expressed genes, with 551 upregulated and 368 downregulated genes, were identified in ARHGAP18 KO cells compared to in WT Jeg-3 cells. The distribution of these DEGs was displayed in the volcano plot (Fig. 3B, Supplemental Data).

**Figure 3:**
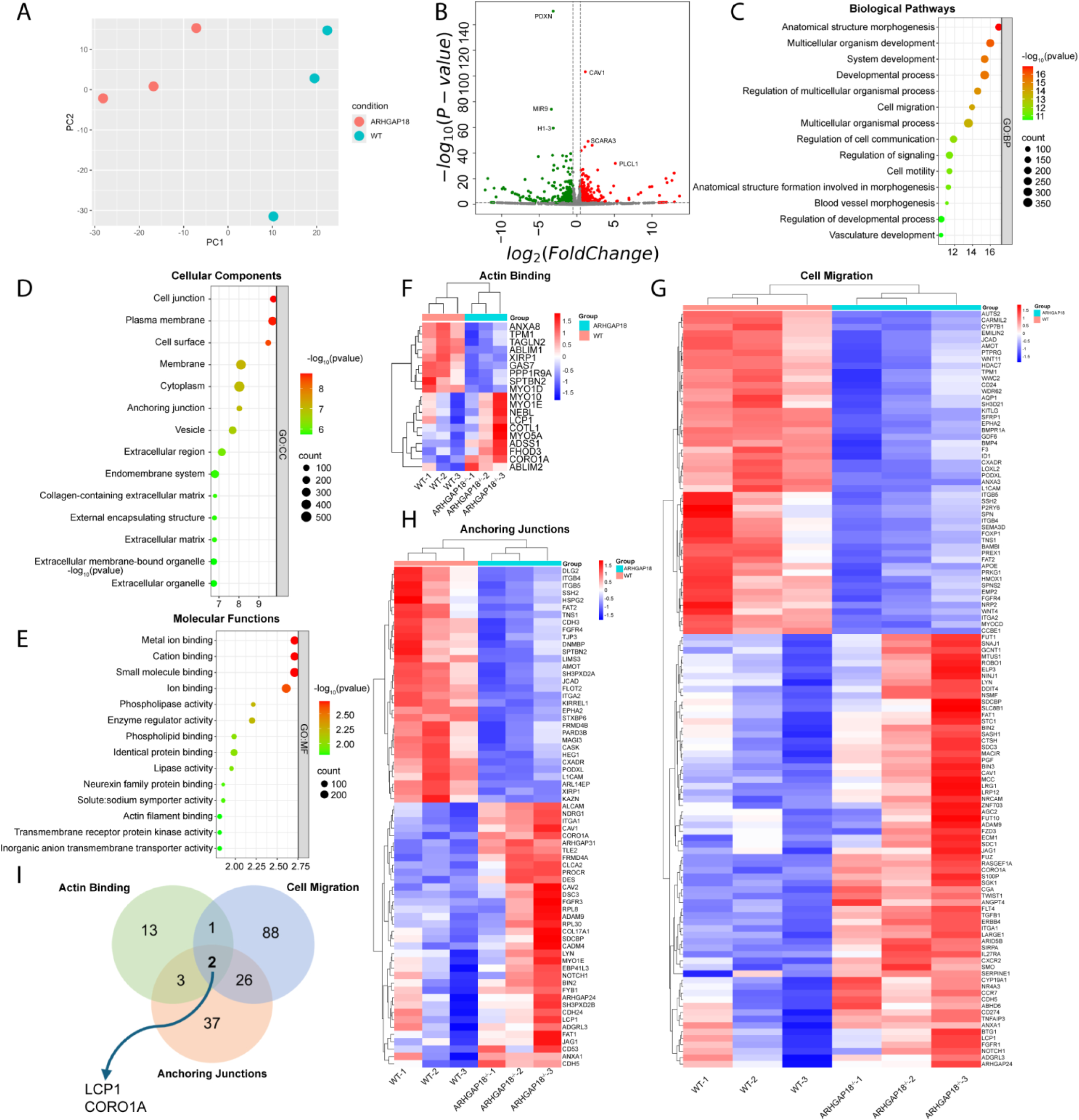
ARHGAP18 total RNA sequencing shows changes in transcription of actin regulatory processes including HIPPO and YAP/TAZ dependent pathways. A) Principle Component Analysis (PCA) from total RNA sequencing results obtained from WT Jeg-3 and ARHGAP18^-/-^ cells (n=3) B) plot of Adjusted p-value vs. log fold change in expression (volcano plot) with select genes identified C) Go terminology analysis of biological pathways. Color indicates p-value, representing the likelihood that a pathway is altered in ARHGAP18^-/-^ vs. WT Jeg-3 cells while bubble size indicates the number of genes identified with altered expression. D) GO terminology analysis of Cellular components E) GO terminology analysis of Molecular Functions. F) Heat map of the genes where the gene product is a known or predicted actin binding protein in each of the 3 biological replicates from WT Jeg-3 or ARHGAP18^-/-^ cells. Color indicates fold change where blue is lower expression and red is increased expression compared to the other group. G) Heatmap similar to (F) for Cell Migration gene products H) Heatmap similar to (F) for Anchoring Junctional gene products. I) Venn diagram identifying genes that are combinations of the terms shown in (F)-(H). LCP1 and CORO1A’s gene products bind actin and have been characterized to play a role at anchoring junctions and in cell migration.

To further explore the impact of ARHGAP18 KO on overall cell functions, we performed Gene Ontology (GO) enrichment analysis and identified that across different GO categories, cell morphology, migration, junctional anchoring, and actin filament binding terms were significantly altered in cells lacking ARHGAP18 (Fig. 3C-E). Additionally, numerous terms related to microvilli and the apical plasma membrane including ion binding, lipid binding, plasma membrane binding were also enriched. To investigate the expression changes of specific genes associated with these terms across all of our samples, we plotted the expressional patterns of each gene into clustered heatmaps (Fig. 3F-H). Microvilli specific regulators included multiple myosin motor proteins responsible for membrane to plasma membrane linkages (Myo1E, Myo1D) and transport of lipid and protein vesicles inside microvilli (Myo10). By comparing the proteins that meet the shared characteristics of anchoring, migration and actin binding (Fig. 3I) we identified LCP1, also known as L-plastin as upregulated in ARHGAP18-deficient cells. Plastin-1 is a predominate actin bundler in microvilli (also called Fimbrin), while L-plastin is associated with tumorigenesis when expressed outside of bone. A second gene, CORO1A (Coronin 1), was also identified, which is involved with RhoA-dependent signaling (Priya et al., 2016). These results are validated by our previous characterization of ARHGAP18 in the biogenesis and maintenance of microvilli at the apical membrane through RhoA (Lombardo et al., 2024).

Further analysis of the individual gene products associated with actin, cell migration, and anchoring junctions identified proteins associated with the defects we previously identified in ARHGAP18 KO cells at the basal surface. Most notable among these were the alterations in the expression of Integrins (ITGA1, ITGA2 INTGB5, INTGB4) and other focal adhesion gene products such as TNS1(Fig. 3F-H). These results are validated by our finding that ARHGAP18^-/-^ cells lack focal adhesions (Fig. 2B). To further investigate the relationship between genes with altered expression in ARHGAP18 KO, we performed network analysis using Ingenuity Pathway Analysis (IPA). Our analysis revealed that some protein-protein relationships can be predicted, such as ARHGAP18’s regulation of RhoA (Fig. S1). However, network analysis also revealed that most interactions would not be through known protein-protein interactions or through RhoA signaling. These results promoted the hypothesis that the alterations defined in the RNAseq data derive from alterations in transcription rather than through direct protein-protein interactions. To explore this potential, we manually characterized the known signaling pathways involved with all down expressed genes related to cell migration (Supplemental Table 1). We found that of these genes, more than half of them were associated with WNT, HIPPO, or MAPK/PI3K signaling pathways, and multiple were direct YAP/TAZ effectors (Supplemental Table 1). We concluded that loss of ARHGAP18 was not only altering cytoskeletal organization through direct RhoA signaling but additionally resulting in alterations to transcriptional signaling of and expression of proteins regulating the actin cytoskeleton.

### ARHGAP18 regulates YAP phosphorylation and nuclear localization

Multiple prior studies indicated that ARHGAP18 interacts downstream of YAP through the Hippo pathway to regulate transcription of actin organization and cell proliferation (Coleman et al., 2020; Porazinski et al., 2015). We hypothesized that the aberrant actin organization observed in ARHGAP18^-/-^ cells, along with the invasive properties of overexpressing cells, resulted from a partial loss of the terminally differentiated apical/basal polarity characteristic of microvilli expressing cells. One possibility was the direct binding of ARHGAP18 to YAP or TAZ. A second possibility was that ARHGAP18 could influence YAP through upstream Hippo pathway components. Our group recently identified a short binding motif of ARHGAP18 from V10 to S17 that bound directly to the FERM domain of ERMs (Lombardo et al 2024). Merlin, which localizes to cell-cell junctions, acts upstream of YAP through the Hippo pathway shares a close structural similarity to ERMs. We tested if ARHGAP18 could form a complex with Merlin using a pull-down approach by passing lysate from WT Jeg-3 cells expressing a tagged Merlin construct over an immobilized ARHGAP18 column. We found that Merlin expressed in human Jeg-3 cells bound to the ARHGAP18 column (Fig. 4A). To investigate this interaction, we sought to identify if ARHGAP18 was localized to junctions where Merlin is active. We used the rescue of ARHGAP18 ^-/-^ cells with a flag tagged ARHGAP18 to image the localization of ARHGAP18 in serum-starved versus cells supplemented with FBS for 1 hour following serum starvation (Fig. 4B). Maximum projection images of ARHGAP18-flag showed localization of ARHGAP18 to microvilli at the apical surface, along with the rescue of increased microvilli formation in ARHGAP18^-/-^ cells as we’ve previously reported (Fig. 4A) (Lombardo et al., 2024). Additionally, ARHGAP18 localized strongly to the nucleus and junctions while being almost entirely excluded from the cytoplasm. Hippo pathway signaling responds to multiple external cues, including nutrient availability to signal downstream transcription for cell proliferation and cytoskeletal remodeling. When cells were deprived of serum, ARHGAP18’s localization shifted to include substantial cytoplasmic staining (Fig. 4B). Interestingly, ARHGAP18’s localization to microvilli and its reduction of the formation of microvilli was maintained in both conditions, indicating that the microvilli specific functions were independent from nutrient availability. We concluded that ARHGAP18 acted in part through a molecular interaction with Merlin at junctions. However, the results also indicated that ARHGAP18 may have a secondary, nuclear and cytoplasmic regulatory function depending on nutrient availability.

**Figure 4:**
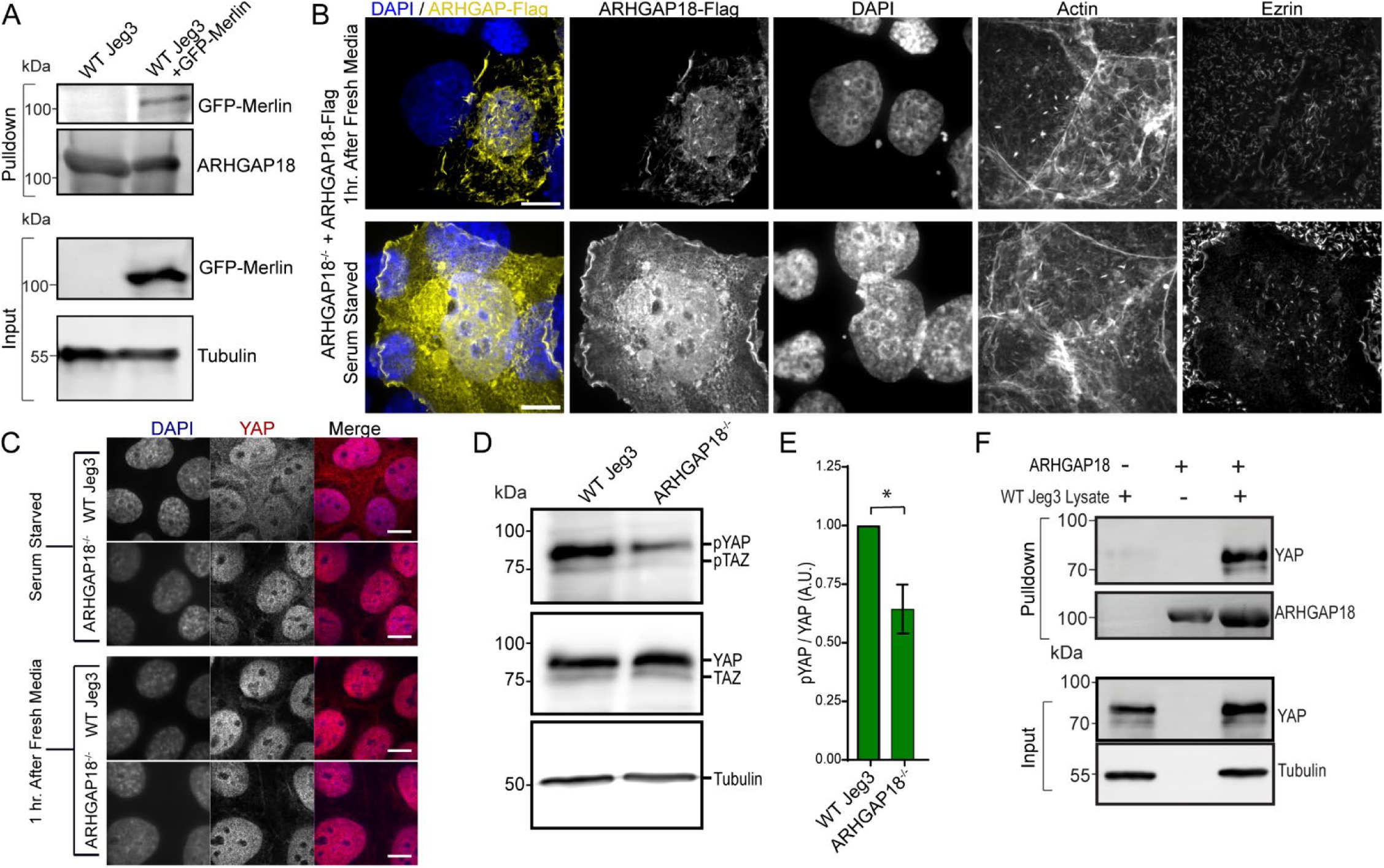
ARHGAP18 Complexes with YAP Affecting its Localization throughout the Cell Based on Nutrient Availability. A) Western blot of an ARHGAP18 pulldown from WT Jeg-3 cells with or without expression of GFP-merlin. B) Immunofluorescent imaging of ARHGAP18-flag rescue of ARHGAP18^-/-^ cells in the serum starved or 1 hour after reintroduction of serum. ARHGAP18’s localization includes the cytoplasm in the serum starved state. Ezrin staining of microvilli shows lower levels of microvilli in both conditions. Scale bar 10µm. C) Immunofluorescent imaging of YAP in WT and ARHGAP18^-/-^ cells in the serum starved state or 1 hour after reintroduction of serum. YAP localization to the cytoplasm during serum starvation is lost in ARHGAP18^-/-^ cells. Scale bar 10µm. D) Western blot of lysate from WT or ARHGAP18^-/-^ cells shows a decrease in the phosphorylation of YAP and TAZ (pYAP/ pTAZ) in the knockout cells. E) Quantification of the band intensity of pYAP/YAP as shown by the blot in (D). Bars indicate Mean±SEM; conditions are significant by paired t-test (p ≤0.05) n=3 F) Pulldown experiment using an ARHGAP18-agarose column. YAP does not bind to the column agarose alone nor is YAP present on the purified ARHGAP18 column before the addition of WT Jeg-3 lysate. When Jeg-3 lysate passed over the ARHGAP18 column endogenous YAP stably binds.

In serum-starved WT cells YAP is phosphorylated and localized to the cytoplasm. Upon reintroduction of nutrients, YAP phosphorylation is reduced, and it is shuttled to the nucleus, where it acts to promote cell proliferation and actin polymerization. Next, we tested if YAP shuttling between the nucleus and cytoplasm in response to nutrient availability was altered in Jeg-3 cells lacking ARHGAP18. We used immunofluorescent staining of YAP and TAZ in cells fixed in either serum-fed or serum-starved states. In WT serum-starved cells, YAP/TAZ showed diffuse staining throughout the cytoplasm and nucleus (Fig. 4C). Upon reintroduction of serum into the media, YAP/TAZ’s cytoplasmic localization was lost within one hour, and only nuclear staining remained. In ARHGAP18 ^-/-^ cells YAP/TAZ remained predominantly in the nucleus in both the serum-starved and serum-present media states. Using lysates from the serum-starved cells, we tested by western blotting the phosphorylation state of YAP and TAZ (Fig. 4D). While the total expression of YAP and TAZ remained the same in WT and ARHGAP18 KO cells, the phosphorylation of YAP and TAZ was significantly reduced in the cells lacking ARHGAP18 compared to WT levels (Fig. 4D, E).

We hypothesized that ARHGAP18 may also form a complex with YAP that influences YAP’s cytoplasmic localization. To test this hypothesis, we passed WT Jeg-3 lysates over the immobilized ARHGAP18 column to probe for binding of native proteins to ARHGAP18. We detected that endogenous YAP from WT Jeg-3 cells bound to immobilized ARHGAP18 (Fig. 4F). We concluded that YAP’s localization to the cytoplasm in the serum starved state depended on the formation of a complex including both YAP and ARHGAP18. The combined results indicate that ARHGAP18 acts as both a negative RhoA regulator and as part of a Hippo signaling feedback loop between YAP and Merlin to control polarized actin cytoskeletal structures.

## Discussion

Humans have 20 Rho/Ras family of GTPase genes where nucleotide state transitions are accelerated by a plethora of guanine nucleotide exchange factors (GEFs) and GTPase Activating Proteins (GAPs). This genetic complexity combined with partial overlapping molecular functions, has presented significant challenges to the characterization of the molecular functions of individual GAPs and GEFs in humans. For example, of the approximately 83 GEFs in humans, at least 24 have specific or partial activity for a single GTPase, RhoA (Vigil et al., 2010). Humans also have genes for three dissociation inhibitors (RhoGDIs), which regulate Rho family localization through membrane binding. Thus, much of our foundational understanding derives from careful experimentation in more genetically advantageous model systems (Etienne-Manneville and Hall, 2002). Here, we use CRISPR KO of human epithelial cells to investigate the loss of a single RhoA effector, ARHGAP18. We uncover that ARHGAP18 forms complexes with YAP and Merlin which drive interwoven molecular pathways regulating actin organization, cell morphology, and junctional integrity. These results indicate that Rho family effectors have the potential to act in complex and multifaceted cellular functions linking multiple signaling and polarity-driving pathways.

We find the naming of the fly ortholog of ARHGAP18 as conundrum (Conu) appropriate (Neisch et al., 2013). Our results add to multiple prior reports indicating that ARHGAP18 has apparent contradictory additional cellular functions outside of Rho GAP activity (Coleman et al., 2020; Lay et al., 2019; Neisch et al., 2013; Porazinski et al., 2015). ARHGAP18 was first identified as a novel Rho effector in a microarray screen of proteins regulating capillary tube formation by human umbilical vein endothelial cells (Coleman et al., 2010). The fly ortholog conundrum (Conu) was subsequently reported to localize to the cell cortex through a molecular interaction with the only fly ERM, Moesin (Neisch et al., 2013). However, increased expression of conu did not produce a phenotype typical of reduced levels of Rho1 as expected from a GAP and instead led to proliferation and cell overgrowth. We identify the same unexpected phenotype in our ARHGAP18 overexpression in human epithelial cells, where cells invade into neighboring monolayers (Fig. 2C).

ARHGAP18^-/-^ cells exhibit a near total loss of stress fibers (Fig. 1E, 2D) despite increased cellular levels of pMLC (Fig. 1B) (Lombardo et al., 2024). These results represent the strongest argument that ARHGAP18 functions in additionally alternative pathways to RhoA/C alone. Our data raises several questions about how RhoA and YAP are coordinated: First, what is the effect of ARHGAP18 complexing with both Merlin and YAP? Loss of ARHGAP18 results in increased active RhoA at junctions (Lombardo et al., 2024) promoting apical actin reorganization. Thus, it’s reasonable to conclude that one of Merlin’s functions as a scaffolding protein is to sequester ARHGAP18 to junctions to regulate RhoA locally (Fig. 5). When proliferative signaling is low, merlin is inactivated and dissociates from the junctional cytoskeletal structures. Inactivation of Merlin would mask the predicted binding site for ARHGAP18 in the conserved FERM domain shared with ERMs. ARHGAP18 would then be free to complex with YAP in the cytosol. The shuttling we characterize of ARHGAP18 localizing from junctions to the cytoplasm in response to serum availability (Fig. 3A) matches this proposed mechanism (Fig.5). This system would place ARHGAP18 as a signaling link between the Hippo and RhoA cascades and as a component in a negative feedback loop between Merlin and YAP.

**Figure 5:**
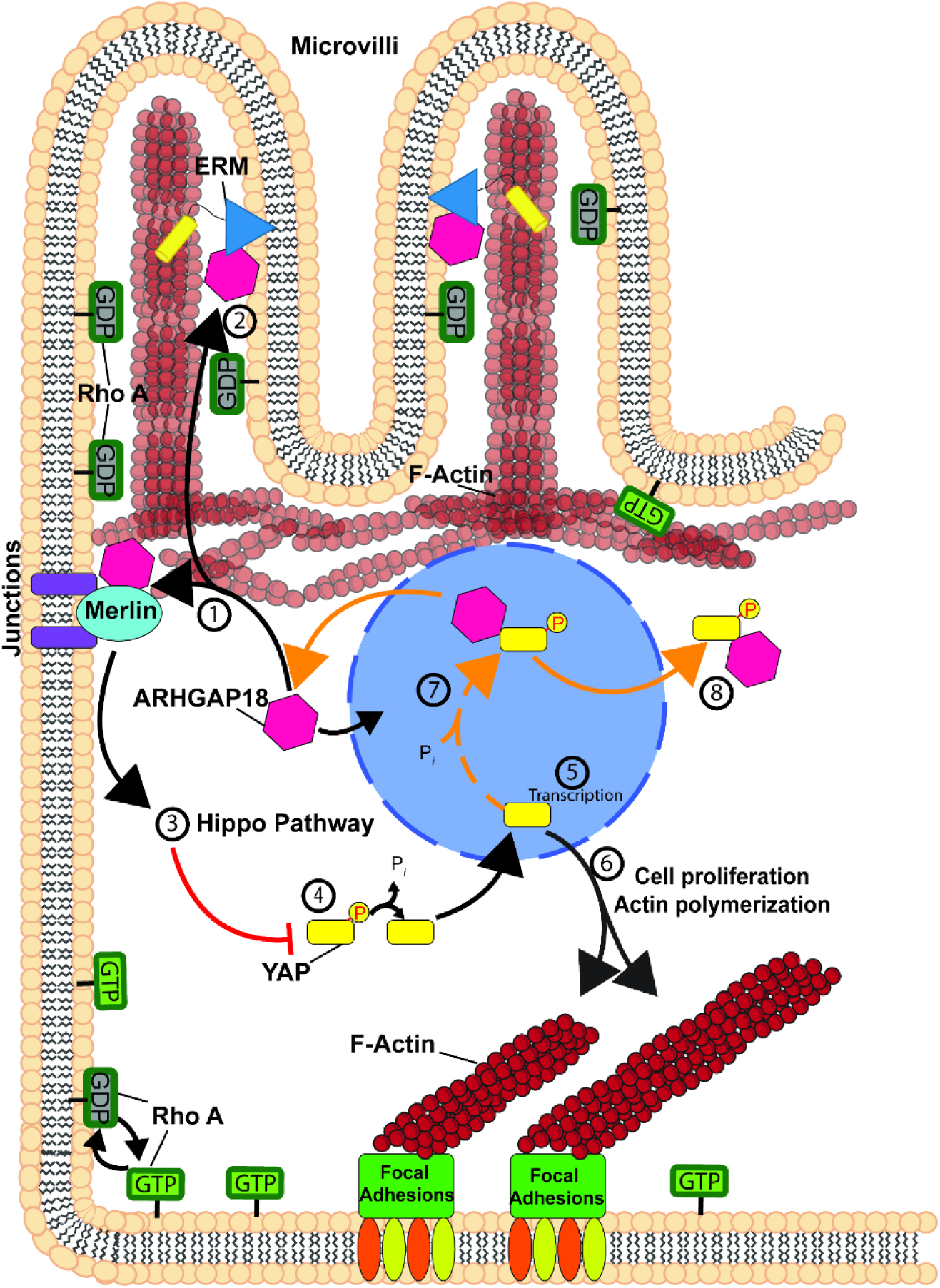
Model of ARHGAP18’s coordination of RhoA and Hippo signaling pathways in Human epithelial cells. Under normal cellular conditions (black arrows), ARHGAP18 localizes to the nucleus, cell junctions, and microvilli. ARHGAP18 forms complexes with (1) Merlin at cell junctions and (2) ERMs in microvilli through direct binding to their conserved FERM domains. Once recruited to junctions by Merlin, ARHGAP18 regulates RhoA locally. Merlin simultaneously coordinates mechanosensation and proliferation signaling through activation of the hippo pathway. (4) Hippo signaling regulates the phosphorylation of YAP, where the active, dephosphorylated state is shuttled to the nucleus. (5) Active YAP/TAZ coordinate transcriptional changes which result in (6) expression of proteins regulating the actin cytoskeleton and proliferation. 7) Alternatively, when Hippo signaling suppresses YAP activation (orange arrows), YAP is phosphorylated through an ARHGAP18 dependent mechanism. 8) ARHGAP18 complexes with YAP which is required for proper phosphorylation and eventual shuttling of YAP to the cytosol for degradation. Loss of ARHGAP18 results in persistent localization of YAP to the nucleus with constitutively active proliferation signaling in addition to misregulation of RhoA at junctional and apical membranes. Without coordination of Hippo and RhoA signaling by ARHGAP18, knockout cells present a loss of stress fibers and filopodia at the basal surface through YAP signaling with disruption of microvilli actin core bundles and junctional integrity at the apical surface through RhoA signaling.

Second, why is non-muscle myosin 2 (NM-2) not producing actin bundles despite being active in cells lacking ARHGAP18? The parameter space for defining this phenotype makes experimentation extremely challenging as actin bundlers, nucleators, severing proteins, and polymerization effectors may all be affected by the alterations to YAP and RhoA in ARHGAP18^-/-^ cells. The list of available mechanisms that YAP alone may induce under perpetual nuclear localization (Fig. 4C) is daunting (Pocaterra et al., 2020). However, our RNAseq analysis narrows the list of potential mechanisms (Fig. 3). One potential explanation, supported by our RNA analysis, is the loss of the mechanical attachment point for the stress fiber bundles at the basal membrane. Integrin expression is severely disrupted in ARHGAP18 deficient cells (Fig. 3F-H) which would deprive a forming actomyosin bundle from mechanically attaching to the plasma membrane. Further insight into the potential mechanism driving the loss of stress fibers could come from predictive modeling using *silico* simulations. These have been productive in defining the minimum components needed to organize complex cytoskeletal behaviors (Landino et al., 2021; Walcott and Warshaw, 2022). This approach will likely be needed in future investigations to predict how alterations in actin filament and myosin dynamics prevent stress fiber formation in ARHGAP18^-/-^ cells. Prior experimental work can also provide some clues to NM-2 overactivation resulting in a loss of actin bundles. In microvilli the loss of ARHGAP18 results in overactivation of NM-2 (specifically isoform MHY14) where motors produce enough force to shred the actin bundles of microvilli (Lombardo et al., 2024). This mechanism occurs to a lesser extent in the apical terminal web of actin in WT epithelial cells through NM-2C (Chinowsky et al., 2020). The force required to dissociate individual actin filaments from a bundle crosslinked by fimbrin (microvilli), fascin (filopodia), or alpha actinin (stress fibers) is unknown and will require careful *in vitro* biophysical assays using atomic force microscopy or optical tweezers to define.

Regardless of the genetic background, a major technical challenge for the study of Rho-dependent signaling pathway’s regulation of cellular actin architecture is that 1) actin is the most abundant protein in the cytosol, 2) actin forms three-dimensional interwoven webs of densely packed networks, 3) filaments can turn over on the scale of 30-60 seconds, and 4) the single filament structure is 7 nanometers across (two orders of magnitude below the diffraction limit of visible light). We’ve overcome these hurdles using super-resolution STORM microscopy (Xu et al., 2012). While multiple prior studies have observed the change in actin bundles of cells with altered ARHGAP18 levels (Coleman et al., 2020; Neisch et al., 2013; Porazinski et al., 2015), our super resolution characterization allows for the dissection of the exact actin structures lost. Reorganization toward branched or individual filaments would dramatically alter the transport of intracellular cargos by myosin and microtubule-based motors (Bensel et al., 2024; Heaslip et al., 2014; Lombardo et al., 2017; Lombardo et al., 2019). Our STORM data indicates that not all actin or bundles are lost as it appears by our own traditional confocal staining (Compare Fig. 2A to Fig.1E). Instead, significant quantities of single filaments are maintained in areas that appear devoid of actin under conventional light microscopy. When these findings are combined with our RNAseq data the molecular pathways responsible for these changes were identified to be a combination of RhoA signaling and pathways both upstream and downstream of YAP/TAZ. Our data indicates that the loss of ARHGAP18 switches the epithelial cell’s basal actin signaling toward a state where the actin architecture resembles a leading edge comprised of single and branching filaments. Consequently, Jeg-3 cells which typically form monolayers, instead form 3-dimensional stacks on top of their neighboring cells when ARHGAP18 signaling is disrupted (Fig. 2C)

Collectively, our results support a model where loss of ARHGAP18 depolarizes cells by dysregulating apical, junctional, and basal membrane identification through ERMs, Merlin, and YAP, respectively. The exact molecular interactors responsible for the signaling cascade that results in pMLC activation in the overexpression of ARHGAP18 will require future characterization. Additional investigation of the molecular details of the ARHGAP18/YAP binding complex will elucidate the mechanism that ARHGAP18 uses to coordinate RhoA signaling with the Hippo pathway to define cell polarity and morphology.

## Methods

### Reagents and cDNAs

Anti-ezrin antibody (CPTC-ezrin-1 supernatant concentrate obtained from the Developmental Studies Hybridoma Bank; catalog no. CPTC-Ezrin-1; Research Resource Identification AB_2100318) was used 1:100 for immunofluorescence. Mouse anti-Flag (Sigma-Aldrich; catalog no. F1804) was used at 1:250 for immunofluorescence and 1:5,000 for Western blot, and mouse anti-tubulin (Sigma-Aldrich; catalog no. T5168) was used at 1:5,000 for Western blot. anti-MLC2 (catalog no. 3672) and anti–phospho-MLC2 (phospho-Thr18/Ser19; catalog no. 3674) were purchased from Cell Signaling Technology and used at 1:500 in Western blots. Anti–nonmuscle myo-IIB from BioLegend (catalog no. 909902) and non-muscle myo-IIA (BioLegend; catalog no. 909802) were used at 1:100 in both Western blots and immunofluorescence. The rabbit ARHGAP18 antibody was produced and characterized previously (Lombardo et al., 2024) and used at 1:500 for western blotting. The rabbit anti-vinculin antibody was a kind gift of A. Bretscher and was described previously (Franck et al., 1990) and used at 1:200 for immunofluorescence. For actin staining, Alexa Fluor 647 phalloidin (Invitrogen; catalog no. A30107) was used at 1:250.

Human ARHGAP18 constructs were produced using polymerase chain reaction (PCR) with New England Biolabs (NEB) Phusion High-Fidelity PCR Kit (Cat# E0553L). cDNA of ARHGAP18 was obtained off a construct originally derived from the Harvard Plasma Database (ID # HsCD00379004). NEB Monarch PCR & DNA Cleanup Kits (Cat# T1030S) and Thermo Scientific GeneJET Gel Extraction Kits (Cat# K0692) were used to purify PCR and Gibson Assembly products. DNA products were cloned into the mammalian expression vector PQCXIP using NEB Gibson Assembly cloning Kit (Cat# E5510S). transformations were done into OneShot TOP10 bacteria (Thermo Fisher; Cat# C404010), selected using ampicillin resistance, and then sequenced for verification. GFP-merlin was a gift of A. Bretscher and produced previously (Sher Bretscher Dev Cell 2012)

### Western Blotting

Western blot analysis of cell lysates was done using 4–12% Thermo Fisher bolt SDS-PAGE gels (Cat# NW04120). Gels were transferred to a polyvinylidene difluoride (PVDF) membrane using a BioRad Transblot Turbo (Cat# 1704150) and blocked with 5% milk in Tris Buffered Saline (TBS) + 0.5% Tween-20 (TBST). For detecting MLC and pMLC, 0.2- µm pore size PVDF (EMD Millipore; Immobilon-PSQ) was used. Primary antibodies were incubated with the membrane in TBST supplemented with 5% BSA either for 1 hr at room temperature or overnight at 4°C. Bands were detected with infrared fluorescent secondary antibodies (Invitrogen or LI-COR Biosciences; catalog nos. 926-32221 and 926-32210). Membranes were imaged using a Bio-Rad MP ChemiDoc. ImageJ intensity profile built-in toolset was used for western blotting quantification, which was then averaged, normalized, and plotted in Microsoft Excel.

### Immunofluorescent Imaging

Room temperature (23°C) fixed cell confocal imaging was done using a spinning-disk (Yokogawa CSU-X1; Intelligent Imaging Innovations) Leica DMi600B microscope with a spherical aberration correction device and either a ×100/1.46 NA Leica objective. Hamamatsu ORCA-Flash 4.0 camera metal-oxide semiconductor device was used to capture images, and Z-slices of acquired images were assembled using SlideBook 6 software (Intelligent Imaging Innovations). Maximum-intensity projections were assembled in SlideBook 6 or ImageJ and exported to Adobe Illustrator for editing. Vertical expansion of side projections was used to increase visual clarity of apical/basal localization. Widefield microscopy was performed using an inverted Leica DMi8 widefield microscope equipped with a Leica ×100 NA air objective, a Leica DFC 9000 GTC camera, Leica Application Suite X, and Leica adaptive focus control.

### Immobilized ARHGAP18 pulldowns

To determine the interaction between ARHGAP18 and Merlin or YAP, pulldown assays were performed using cell lysate from Jeg-3 WT cells. Purified human ARHGAP18 was produced by bacterial expression of an N-terminal-SUMO-HIS-tagged protein purified using a NiNTA resin as described in (Lombardo et al., 2024). Transfections for GFP Merlin were performed using PEI MAX polyethylenimine reagent (Polysciences; Cat# 24765). Cells were harvested with lysis buffer (25 mM Tris, 5% glycerol, 150 mM NaCl, 50 mM NaF, 0.1 mM Na3VO4, 10mM βGP, 0.2% Triton X-100, 250 mM calyculin A, 1 mM DTT, 1× cOmplete Protease Inhibitor Cocktail [Roche; Cat# 11836153001]) by scraping. Lysates were then sonicated, and centrifuged at 8000 × g for 10 min at 4°C. Before incubating with the cell lysates, SUMO-ARHGAP18 NiNTA beads were equilibrated and washed into lysis buffer. The sample of the supernatant was taken for input, then the rest was added to the SUMO-ARHGAP18 NiNTA beads and nutated for 3 hr at 4°C. After incubation, the beads were pelleted and washed four times with a 2 minute incubation before boiling in 40 µL 2× Laemmli sample buffer.

### RNA-Seq Sample Preparation

Cells were plated in 100mm Corning 100 × 20 mm Petri-style TC-treated culture dishes (Cat# 430167) and allowed to grow to 80% confluency. Cells were plated at 10am on day 0, media was exchanged at 10am on day 1 and cells were collected and pelleted at 10am on day 2 over a total of 48 hours to maintain environmental consistency. Pelleting of cells was performed by washing the cells 3 times in PBS then adding 0.05% trypsin (Thermo Fisher Scientific; Cat# 25300054) and incubating at 37°C and 5% CO2 for 5 minutes. Once detached cells were washed into MEM media (Thermo Fisher Scientific; Cat# 10370088) then the media was exchanged for ice cold PBS. Cells were pelleted at 1000x g and stored at −80C until they were lysed for RNA extraction. Total RNA was extracted at the University at Buffalo Core Genomics Facility using TRIzol reagent (cat. no. 15596018, Invitrogen). It was then purified with an RNeasy kit (cat. no. 74106, Qiagen). Libraries were prepared using the Illumina TruSeq Stranded total RNA kit and the Ribo-Zero plus rRNA depletion kit. The sequencing was performed on an Illumina HiSeq 4000 PE100 sequencer. The nf-core/rnaseq Version 3.18.0 bioinformatics pipeline was used to analyze RNA sequencing data with alignment and quantification performed using STAR (Dobin et al., 2013) followed by Salmon (Patro et al., 2017).

### Bioinformatic analysis

Raw gene counts were analyzed using DESeq2 package in R to identify differentially expressed genes (DEGs) between WT-Jeg3 and ARHGAP18 KO conditions. Genes were classified as DEGs if the following filtering criteria were met: false-discovery rate (FDR) adjusted p-value of < 0.05 and absolute log_2_(fold-change) of > 0.5. Sample distribution was visualized with the Principal Component Analysis (PCA) plot generate by ggplot2 package in R using a normalized count dataset. Python’s bioinfokit was used to create the volcano plot.

Functional enrichment analysis was performed using the g:GOSt function on the gProfiler web server (https:// biit.cs.ut.ee/gprofiler/gost) with the identified DEGs. The significance threshold was set to FDR value, and significant Gene Ontology (GO) terms in molecular functions, biological pathways, and cellular components categories were defined by an adjusted p-value of < 0.05. The top 15 GO terms in each category were visualized using bubble plots generated by SRplot online server.

To visualize expression patterns across samples, clustered heatmap was generated with SRplot online server using log_2_-transformed normalized gene, expression values. Specifically, hierarchical clustering was performed on both genes (rows) and samples (columns) using Euclidean distance and complete linkage cluster methods. Genes visualized were DEGs annotated in the GO term.

Ingenuity pathway analysis (IPA, QIAGEN) was used to perform network analysis on 889 DEGs identified in this dataset. A core analysis was run on this dataset, which produced information on various mechanistic pathways and enriched function-based literature gathered in IPA knowledge base. Specifically, we used the “My Pathway” function to generate predicted networks between ARHGAP18 and some genes of interest. The “Molecular Activity Predictor” tool was used to elucidate predicted activation states of molecules and interactions downstream of ARHGAP18.

### Cell culture and expression vectors

Jeg3 cells were purchased from ATCC.org (Cat# Htb-36) and maintained in a humidified incubator at 37°C and 5% CO2. Media for Jeg3 cells used in 1× MEM (Thermo Fisher Scientific; Cat# 10370088) with penicillin/streptomycin (Thermo Fisher Scientific; Cat# 15070063), 10% fetal bovine serum (FBS; Thermo Fisher Scientific; Cat# 26140079), and GlutaMAX (Thermo Fisher Scientific; Cat# 35050061). Corning 100 × 20 mm Petri-style TC-treated culture dishes (Cat# 430167) were used for maintaining cultures. Transient transfections used a PEI MAX polyethylenimine reagent (Polysciences; Cat# 24765). All cell lines were checked for mycoplasma in regular intervals using DAPI staining. ARHGAP18 knockout cells were created and characterized (Lombardo et al., 2024). In summary of these methods, multiple CRISPR single-guide RNAs against the sequence 5′-CAGCGGCAAGGACCAGACCG-3′ or 5′-CAGCGGCAAGGACCAGACCG-3′ were cloned into puromycin-resistant pLenti-CRISPRV2 (Addgene; Cat# 49535) which was transfected into 293TN cells with psPAX3 and pCMV-VSV-G (a gift from Jan Lammerding, Weill Institute for Cell and Molecular Biology, Cornell University, Ithaca, NY) for 48–72 hr. Jeg3 cells were then transduced with virus supplemented with Polybrene to 8 µg/mL twice a day for 2 days, followed by puromycin selection at 2 µg/mL. Mixed populations of selected cells were single-cell sorted and then screened by western blotting. ARHGAP18 expression vectors were created and described in (Lombardo et al., 2024).

### Super-Resolution STORM

Jeg-3 cells were plated on 35 mm 1.5 high precision glass bottom MatTek dishes (Cat# P35G-0.170-14-C) and fixed and permeabilized using 2mLs of 0.3% glutaraldehyde and 0.25% Triton X-100 in cytoskeleton buffer (10 mM MES pH 6.1, 150 mM NaCl, 5 mM EGTA, 5 mM glucose, and 5 mM MgCl_2_) for 1-2 min at room temperature. Initial fixation was followed by 2% glutaraldehyde in cytoskeletal buffer for 10 min at room temperature. The sample was then treated with 1mL 0.1% NaBH4 in Phosphate Buffered Saline (PBS) at pH 7.4 for 7 min to reduce background fluorescence. Actin was stained in phalloidin at a dilution of 1:250 overnight at 4°C. Slides were washed into 50mM Tris, pH 8.0, 10% glucose, 10mM NaCl, 0.5mg/ml Glucose oxidase, 0.04 mg/ml catalase with 10% cysteamine by weight, and 1.5% 2-Mercaptoethanol (BME) by weight. Stochastic Optical Reconstruction Microscopy (STORM)(Rust et al., 2006) was performed using a Zeiss Elyra super-resolution inverted Axio Observer.Z1 microscope provided by Cornell University’s Institute of Biotechnology Imaging Facility. Lasers emitting 405 and 640 nm wavelengths through a 100x/1.46 NA oil objective captured images on a pco.edge 5.5m camera. Exposure time ranged from 50 to 400 ms and was dependent on the sample and channel to optimize STORM reconstruction. A minimum of 100,000 images were used for each STORM processing using the Imagej ThunderSTORM toolset (Ovesny et al., 2014) or The Zeiss Zen built-in STORM reconstruction toolset and adjusting the filtering conditions to maximize the signal to noise of single actin filaments in reconstructions.

### Statistical Methods

Statistical comparisons were performed in Microsoft Excel. The type of statistical tests utilized, the configuration of error bars, and the number of independent data points (*n*) are detailed in the figure legends respective to the data tested. Nonparametric or parametric testing was justified through the assumption of the tested data to have a normal distribution or not. Analyses were also done using GraphPad Prism to produce a graphical representation of the amount of phosphorylated YAP over the total expression of YAP with the test and *n* listed in the figure legend.

## Acknowledgements

We thank A. Bretscher and R. Visawantha for their kind gift of the human ARHGAP18 plasmid. We thank A. Martin for their critical comments on the data interpretation. Funding was supported by the National Institutes of Health grant R35GM156870 to A.T.L. and grant R01HL163168 to Y.B.. Preliminary data was collected under NIH grant R35GM131751 to A. Bretscher. We thank A. Heaslip and N. Bouffard for their expertise in troubleshooting STORM. Super resolution microscopy was performed at both the Microscopy Imaging Center at the University of Vermont (RRID# SCR_018821), and the Cornell Institute of Biotechnology Imaging Facility supported by the National Science Foundation funding (1428922).

