## Supplemental Table 1 for "The Rho effector ARHGAP18 coordinates a Hippo pathway feedback loop through YAP and Merlin to regulate the cytoskeleton and epithelial cell polarity"

| Gene | Description | Comment | Citation |
|---|---|---|---|
| AUTS2 | Activator Of Transcription And Developmental Regulator AUTS2 | Part of the PRC1-like complex | https://pubmed.ncbi.nlm.nih.gov/25519132/ |
| CARMIL2 | Capping Protein Regulator And Myosin 1 Linker 2 | Membrane bound actin capping protein | https://pubmed.ncbi.nlm.nih.gov/26578515/ |
| CYP7B1 | Cytochrome P450 Family 7 Subfamily B Member 1 | Metabolistm of steroids | https://pubmed.ncbi.nlm.nih.gov/10588945/ |
| EMILIN2 | Elastin Microfibril Interfacer 2 | Microfibrillar Protein | https://pubmed.ncbi.nlm.nih.gov/26945878/ |
| JCAD | Junctional Cadherin 5 Associated Protein | Cell-to-cell junction protein regulating Hippo signaling | https://pubmed.ncbi.nlm.nih.gov/29794114/ |
| AMOT | Angiomotin | Repressor of YAP1 | https://pubmed.ncbi.nlm.nih.gov/21205866/ |
| PTPRG | Protein Tyrosine Phosphatase Receptor Type G | Growth factor receptor | https://pubmed.ncbi.nlm.nih.gov/16263724/ |
| WNT11 | Wnt Family Member 11 | WNT gene family member | https://pubmed.ncbi.nlm.nih.gov/11712081/ |
| HDAC7 | Histone Deacetylase 7 | Regulates activity of FOXP3 | https://pubmed.ncbi.nlm.nih.gov/17360565/ |
| TPM1 | Tropomyosin 1 | Actin binding protein which regulates myosin binding activity | https://pubmed.ncbi.nlm.nih.gov/38971309/ |
| WWC2 | WW And C2 Domain Containing 2 | Hippo Pathway component. Regulates phosphorylation of YAP1 | https://pubmed.ncbi.nlm.nih.gov/24682284 |
| CD24 | Precurer protein to a sialoglycoprotein | Expressed on cell exterior | https://pubmed.ncbi.nlm.nih.gov/32790899/ |
| WDR62 | WD Repeat Domain 62 | MAPK binding protein 1 paralog | https://pubmed.ncbi.nlm.nih.gov/19910486/ |
| AQP1 | Aquaporin 1 | Water channel membrane protein | https://pubmed.ncbi.nlm.nih.gov/9177353/ |
| SH3D21 | SH3 Domain Containing 21 | EGFR signaling | https://pubmed.ncbi.nlm.nih.gov/20029029 |
| KITLG | KIT Ligand | Stimulates phosphorylation of PIK3 and activation of AKT1 with activation of MAP kinases | https://pubmed.ncbi.nlm.nih.gov/32463597/ |
| SFRP1 | Secreted Frizzled Related Protein 1 | WNT Antagonist | https://pubmed.ncbi.nlm.nih.gov/14871816/ |
| EPHA2 | Ephrin Receptor A2 | Inhibition of the ERK1/ERK2 (MAPK3/MAPK1) signaling pathway | https://pubmed.ncbi.nlm.nih.gov/12400011/ |
| BMPR1A | Bone Morphogenetic Protein Receptor Type 1A | Transmembrane serine/threonine kinase invovled in PI3K/AKT signaling | https://pubmed.ncbi.nlm.nih.gov/25110865/ |
| GDF6 | Growth Differentiation Factor 6 | Secreted ligand of the TGF-beta | https://pubmed.ncbi.nlm.nih.gov/23307924/ |
| BMP4 | Bone Morphogenetic Protein 4 | Ligand activating ERK/MAP kinase, PI3K/Akt | https://pubmed.ncbi.nlm.nih.gov/31363885 |
| F3 | Coagulation Factor III | Cell surface glycoprotein initiates the blood coagulation cascade | https://pubmed.ncbi.nlm.nih.gov/25535411/ |
| ID1 | Inhibitor of DNA Binding 1 | Interferes with DNA binding of Transcription factors | https://pubmed.ncbi.nlm.nih.gov/10537105/ |
| CXADR | CXADR Ig-Like Cell Adhesion Molecule | Activates PI3-kinase and MAP kinases | https://pubmed.ncbi.nlm.nih.gov/38146657 |
| LOXL2 | Lysyl Oxidase Like 2 | Transcription corepressor | https://pubmed.ncbi.nlm.nih.gov/27735137/ |
| PODXL | Podocalyxin Like | Ezrin Binding. MAPK / PI3K | https://pubmed.ncbi.nlm.nih.gov/17616675 |
| ANXA3 | Annexin A3 | Negative regulator of the mitogen-activated protein kinase (MAPK) pathway, promoting the targeting of EGFR to lysosomes for degradation | https://pubmed.ncbi.nlm.nih.gov/22797061/ |
| L1CAM | L1 Cell Adhesion Molecule | Regulates cell adhesion and the generation of transmembrane signals | https://pubmed.ncbi.nlm.nih.gov/22796939/ |
| ITGB5 | Integrin Subunit Beta 5 | Integrin regulated by Hippo signaling and YAP/TAZ | https://pubmed.ncbi.nlm.nih.gov/28504269/ |
| SSH2 | Slingshot Protein Phosphatase 2 | Cofillin Hippo | https://pubmed.ncbi.nlm.nih.gov/25864508/ |
| P2RY6 | Pyrimidinergic Receptor P2Y6 | G-Protein Coupled receptor activated by extracellular nucleotides | https://pubmed.ncbi.nlm.nih.gov/39322240/ |
| SPN | Sialophorin (CD43) | ERM binding protein | https://pubmed.ncbi.nlm.nih.gov/10385528/ |
| ITGB4 | Integrin Subunit Beta 4 | Integrin regulated by Hippo signaling and YAP/TAZ | https://pubmed.ncbi.nlm.nih.gov/28504269/ |
| SEMA3D | Semaphorin 3D | Activates Hippo Signaling | https://pubmed.ncbi.nlm.nih.gov/35613278/ |
| FOXP1 | Forkhead Box P1 | Direct YAP/TAZ binding protein | https://pubmed.ncbi.nlm.nih.gov/24525530/ |
| TNS1 | Tensin1 | Focal adhesion protein. Activates YAP. | https://pubmed.ncbi.nlm.nih.gov/38297127/ |
| BAMBI | BMP and activin membrane-bound inhibitor | Target gene of Bmp-4 signaling. | https://pubmed.ncbi.nlm.nih.gov/11165491/ |
| PREX1 | Phosphatidylinositol-3,4,5-Trisphosphate Dependent Rac Exchange Factor 1 | Rac1 GEF stimulated by PI3K/ ERK signaling | https://pubmed.ncbi.nlm.nih.gov/17308088/ |
| FAT2 | FAT Atypical Cadherin 2 | Member of the cadherin superfamily. Supresses YAP activation | https://pubmed.ncbi.nlm.nih.gov/29985391/ |
| APOE | Apolipoprotein E | Activates MAP3K12 and a non-canonical MAPK signal transduction pathway | https://pubmed.ncbi.nlm.nih.gov/28111074 |
| PRKG1 | Protein Kinase CGMP-Dependent 1 | Activates MAPK Kinase (MEK) | https://pubmed.ncbi.nlm.nih.gov/10567406/ |
| HMOX1 | Heme Oxygenase 1 | Catalyzes the oxidative cleavage of heme | https://pubmed.ncbi.nlm.nih.gov/11121422/ |
| SPNS2 | SPNS Lysolipid Transporter 2, Sphingosine-1-Phosphate | Lipid transporter of sphingosine 1-phosphate | https://pubmed.ncbi.nlm.nih.gov/19074308/ |
| EMP2 | Epithelial Membrane Protein 2 | Regulates integrins | https://pubmed.ncbi.nlm.nih.gov/16216233 |
| FGFR4 | Fibroblast Growth Factor Receptor 4 | Cell surface receptor that mediates activation of MAP kinase signaling pathway | https://pubmed.ncbi.nlm.nih.gov/21203561/ |
| NRP2 | Neuropilin-2 | Transmembrane receptor. Binds Semaporin 3 | https://pubmed.ncbi.nlm.nih.gov/28843905/ |
| WNT4 | Wnt Family Member 4 | Wnt Family Member 4 | https://pubmed.ncbi.nlm.nih.gov/24964196/ |
| ITGA2 | Integrin Subunit Alpha 2 | Integrin regulated by Hippo signaling and YAP/TAZ | https://pubmed.ncbi.nlm.nih.gov/28504269/ |
| MYOCD | Myocardin | Regulated by YAP/TAZ | https://pubmed.ncbi.nlm.nih.gov/37927241/ |
| CCBE1 | Collagen And Calcium Binding EGF Domains 1 | Involved in binding to components of the extracellular matrix | https://pubmed.ncbi.nlm.nih.gov/21778431/ |
