## Supplemental Figure 1 for "The Rho effector ARHGAP18 coordinates a Hippo pathway feedback loop through YAP and Merlin to regulate the cytoskeleton and epithelial cell polarity"

A

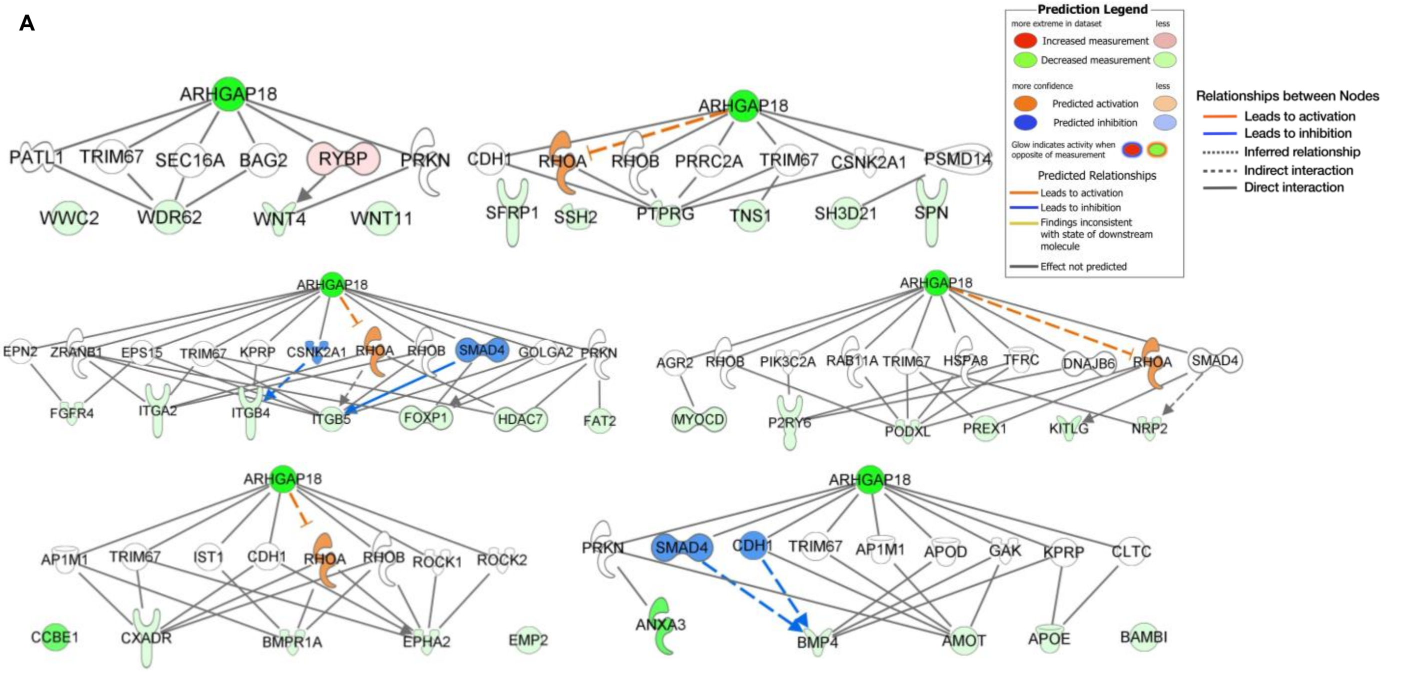


B

### Upstream regulator predicted to be involved due to expressional changes seen in the dataset (predicted inhibition)

| Upstream Regulator | Molecule Type | Predicted Activation State | Activation z-score | p-value of overlap | Target Molecules in Dataset |
|---|---|---|---|---|---|
| FOXA1 | transcription regulator | Inhibited | -3.116 | 5.59E-11 | ↑ANXA1, ↓AQP1, ↑ARHGEF9, ↑BHLHE40, ↑BNIP3L, ↑B... ...all 30 |
| SFTPA1 | other | Inhibited | -3.000 | 3.63E-04 | ↓AMOT, ↑CRLF1, ↑ECM1, ↓GABRP, ↑PLIN2, ↑SERPINE1, ...all 9 |
| N-[N-(3,5-difluorophenacetyl-L-Ala)]-S | chemical - protease inhibitor | Inhibited | -2.944 | 1.91E-03 | ↓CD24, ↑FSTL3, ↑HAPLN1, ↑HEY1, ↑JAG1, ↑LOC10272... ...all 12 |
| miR-141-3p (and other miRNAs w/see | mature microRNA | Inhibited | -2.612 | 1.09E-04 | ↑CD274, ↑JAG1, ↑PITX1, ↑PMAIP1, ↓RHPN2, ↑RNF128, ...all 10 |
| NR0B2 | ligand-dependent nuclear receptor | Inhibited | -2.599 | 1.05E-01 | ↑CARD11, ↓CYP7B1, ↑FGFR1, ↑MAP3K8, ↑PIK3C2G, ↑S... ...all 8 |
| ETV6-RUNX1 | fusion gene/product | Inhibited | -2.571 | 2.25E-03 | ↓ABLIM1, ↓ABR, ↑ANTXR2, ↑ARHGAP24, ↑CARD11, ↑... ...all 19 |
| 26S PROTEASOME (complex) | complex | Inhibited | -2.530 | 1.28E-02 | ↑ATP6V1H, ↑CTSB, ↑CTSF, ↓FOXO1, ↑HSPB8, ↑NOTCH1 ...all 10 |
| CAPN6 | peptidase | Inhibited | -2.530 | 3.76E-04 | ↓BMP4, ↓BMPR1A, ↓FZD7, ↓ID1, ↑SERPINE1, ↓TCF7, ↑.. ...all 10 |
| EGLN (family) | group | Inhibited | -2.459 | 9.06E-04 | ↑BNIP3, ↓DMKN, ↑FSTL3, ↑MAP3K8, ↓MYOCD, ↑PDG... ...all 13 |
| salirasib | chemical drug | Inhibited | -2.433 | 6.87E-03 | ↑BHLHE40, ↑BNIP3, ↑ENO2, ↑PFKFB4, ↑SERPINE1, ↑STC1 ...all 6 |
| CCN5 | growth factor | Inhibited | -2.401 | 4.44E-03 | ↓CD24, ↑PROCR, ↑SERPINE1, ↑TGFB1, ↑TWIST1, ↑ZNF4... ...all 6 |
| miR-155-5p (miRNAs w/seed UAAUGC | mature microRNA | Inhibited | -2.359 | 7.68E-02 | ↑AMIGO2, ↑MYO10, ↑MYO1E, ↑PICALM, ↑PMAIP1, ↓P... ...all 9 |
| ALPHA CATENIN (family) | group | Inhibited | -2.343 | 8.80E-03 | ↑CTSB, ↑ELK3, ↓IGF2, ↑LYN, ↑SGK1, ↑SNAI1, ↑TNFAIP3, ...all 9 |
| ESR1 | ligand-dependent nuclear receptor | Inhibited | -2.254 | 7.16E-10 | ↓ABCA4, ↓ABLIM1, ↑ADCY1, ↑ALCAM, ↓ANGPTL2, ↓A... ...all 96 |
| ibrutinib | chemical drug | Inhibited | -2.236 | 4.51E-01 | ↓BMP4, ↑COX6C, ↓MGLL, ↑RIPK2, ↑WDR74 ...all 5 |
| CELF1 | translation regulator | Inhibited | -2.236 | 6.33E-03 | ↓ADGRL2, ↓DMPK, ↓DNMBP, ↑JAG1, ↓PXDN, ↑WFS1 ...all 6 |
| OVOL2 | transcription regulator | Inhibited | -2.224 | 1.42E-02 | ↑ARHGAP31, ↑FAM13A, ↑NR2F1, ↑SDCBP, ↑WIPF1 ...all 5 |
| TNFSF12 | cytokine | Inhibited | -2.216 | 3.65E-01 | ↑HEY1, ↑HEYL, ↓ID1, ↑MYOD1, ↑NOTCH1 ...all 5 |
| mir-34 (includes others) | microRNA | Inhibited | -2.207 | 2.38E-02 | ↑BNIP3L, ↑FGFR1, ↑HEY1, ↑KLF10, ↓LIN28A, ↑NOTCH1, ...all 8 |
| GIP | other | Inhibited | -2.200 | 2.90E-02 | ↑IKBKB, ↓MX1, ↑SERPINE1, ↓TCF7, ↑TNFRSF1B ...all 5 |
| DCAF1 | kinase | Inhibited | -2.177 | 5.57E-03 | ↑CD274, ↑KLF10, ↑PMAIP1, ↑SOCS2, ↑TNFSF10 ...all 5 |
| PDLIM2 | other | Inhibited | -2.111 | 8.22E-05 | ↑CRISPLD2, ↓DPF3, ↑MYL9, ↑NR2F1, ↑PGCKA1, ↑RAS... ...all 11 |
| iron | chemical - endogenous mammalian | Inhibited | -2.085 | 2.07E-02 | ↑GADD45G, ↓HMOX1, ↓ID1, ↑NDRG1, ↑SLC40A1 ...all 5 |
| IL10RA | transmembrane receptor | Inhibited | -2.065 | 2.02E-03 | ↓ADGRL2, ↓ALDH2, ↑ASS1, ↓BMP4, ↑BNIP3, ↑CCR7, ↓... ...all 19 |
| mifepristone | chemical drug | Inhibited | -2.022 | 2.03E-04 | ↓ACAN, ↑ANXA1, ↓APOE, ↓AQP1, ↑CAV1, ↓CD24, ↑C... ...all 23 |
| N-[(2Z)-3-(4,5-dihydro-1,3-thiazol-2-yl) | chemical reagent | Inhibited | -2.008 | 2.68E-03 | ↑CYP19A1, ↑ELOVL3, ↑LYPD3, ↑NR4A3, ↑SERPINE1, ↑S... ...all 8 |
| HSP70 (family) | group | Inhibited | -2.000 | 8.08E-02 | ↓ACSL3, ↑GBP1, ↑MAPT, ↑SAT1, ↑SLC40A1 ...all 5 |
| LAS1L | other | Inhibited | -2.000 | 9.85E-02 | ↑ASS1, ↑RN7SK, ↑SCARNA13, ↑SMPDL3B ...all 4 |
| SKIC2 | enzyme | Inhibited | -2.000 | 4.28E-02 | ↑BHLHE40, ↑CTH, ↑DDIT4, ↑STC1 ...all 4 |
| ILKAP | phosphatase | Inhibited | -2.000 | 3.89E-03 | ↑CCR7, ↓FOXO1, ↑NDRG1, ↑NR4A3 ...all 4 |

C

**Upstream regulator predicted to be involved due to expressional changes seen in the dataset (predicted activation)**

| Upstream Regulator | Molecule Type | Predicted Activation State | Activation z-score | p-value of overlap | Target Molecules in Dataset |
|---|---|---|---|---|---|
| TP53 | transcription regulator | Activated | 4.101 | 8.87E-10 | ↓ACSL3, ↓ACSL6, ↑AGO2, ↑ANXA1, ↓ANXA3, ↑APOBE.....all 125 |
| dimethyl n-oxalyl-glycine | chemical reagent | Activated | 3.969 | 4.74E-10 | ↑BNIP3, ↑BNIP3L, ↑BTG1, ↑CAV1, ↑CORO1A, ↑DDIT4, ↑.. ...all 28 |
| BHLHE40 | transcription regulator | Activated | 3.797 | 2.03E-05 | ↑ANXA1, ↑BHLHE40, ↑BHLHE41, ↑BTG1, ↑CAV1, ↑CCR7, ...all 25 |
| CD3 (complex) | complex | Activated | 3.557 | 1.13E-05 | ↑ANXA1, ↓ATP11B, ↓BAMBI, ↑BTG1, ↑CD274, ↑CD53, ↑.. ...all 39 |
| IL13 | cytokine | Activated | 3.394 | 2.35E-04 | ↑AVPI1, ↑CISH, ↑CTSB, ↑CTSH, ↑CXCR2, ↑ELP3, ↑ERBB4, ...all 30 |
| CALCA | other | Activated | 3.301 | 3.38E-03 | ↑GAST, ↓HMOX1, ↑IL2RB, ↑JAG1, ↑PLEKHA6, ↑PLXDC2, ...all 13 |
| doxorubicin | chemical drug | Activated | 3.300 | 1.82E-03 | ↑ANXA1, ↑BHLHE40, ↓BMP4, ↑BNIP3L, ↑BTG1, ↑CD274, ...all 40 |
| ZBTB16 | transcription regulator | Activated | 3.293 | 5.87E-02 | ↓AGPAT4, ↓CD24, ↑CD274, ↑CSGALNACT1, ↑DDIT4, ↑F... ...all 11 |
| IL6 | cytokine | Activated | 3.199 | 4.92E-05 | ↓ADGRL2, ↑ANXA1, ↓APOE, ↓BAMBI, ↑BHLHE40, ↑CCR7 ...all 47 |
| lipopolysaccharide | chemical drug | Activated | 3.180 | 4.76E-08 | ↓ACAN, ↑ALCAM, ↓ALDH2, ↓ANGPTL2, ↑ANXA1, ↓A... ...all 138 |
| PIWIL4 | other | Activated | 3.162 | 7.04E-03 | ↑ADSS1, ↑BTG1, ↑ELK3, ↑FYB1, ↑MFGE8, ↑NDRG1, ↑S... ...all 10 |
| PTGS2 | enzyme | Activated | 3.076 | 5.53E-03 | ↑ADGRL3, ↑ANXA1, ↓AQP1, ↑CCR7, ↑CD274, ↑CXCR2, ...all 15 |
| CSF2 | cytokine | Activated | 2.986 | 1.47E-02 | ↑ANXA1, ↓BMPR1A, ↑CARD11, ↑CCR7, ↓CD24, ↑CD274, ...all 25 |
| etoposide | chemical drug | Activated | 2.914 | 2.94E-02 | ↑CD274, ↑CLCA2, ↑CSF2RB, ↑DDIT4, ↑GADD45G, ↑GBP2, ...all 15 |
| olaparib | chemical drug | Activated | 2.897 | 1.24E-04 | ↑BHLHE40, ↑BOK, ↑BTG1, ↑GADD45G, ↑GBP2, ↑GDF15, ...all 14 |
| PDGF-BB (complex) | complex | Activated | 2.830 | 1.15E-04 | ↑BHLHE40, ↓BMP4, ↑CAV2, ↑CTH, ↓F3, ↑FGFR1, ↑GBP2, ...all 24 |
| PD173074 | chemical reagent | Activated | 2.828 | 6.74E-11 | ↑CCR7, ↑CDH5, ↑CGA, ↑CYP19A1, ↑ERVFRD-1, ↓F3, ↓... ...all 18 |
| CD28 | transmembrane receptor | Activated | 2.828 | 1.99E-05 | ↑ANXA1, ↓ATP11B, ↑BHLHE40, ↑BTG1, ↑CD274, ↑CISH, ...all 26 |
| EGF | growth factor | Activated | 2.807 | 6.40E-06 | ↓ACSL3, ↑ALCAM, ↑CAV1, ↑CD274, ↑CDH5, ↑CGA, ↑CI... ...all 44 |
| NLRP7 | other | Activated | 2.795 | 4.47E-07 | ↑APOBEC3C, ↑CD274, ↓FBXO38, ↑FXYD3, ↓MAGEA4, ↑N... ...all 8 |
| tetradecanoylphorbol acetate | chemical drug | Activated | 2.784 | 3.84E-09 | ↓ACAN, ↑ADAM9, ↓ADCY3, ↑ALCAM, ↑AMIGO2, ↑ANX... ...all 77 |
| STAT4 | transcription regulator | Activated | 2.783 | 1.41E-05 | ↑BHLHE40, ↑ENO2, ↑FYB1, ↓FZD7, ↑GADD45G, ↑MAG... ...all 20 |
| AHR | ligand-dependent nuclear receptor | Activated | 2.762 | 4.31E-05 | ↑ADAM9, ↑AHRR, ↓ALDH2, ↑AMIGO2, ↓BAMBI, ↑CAD... ...all 36 |
| 3,5-dihydroxyphenylglycine | chemical reagent | Activated | 2.714 | 3.23E-03 | ↑BCAS1, ↓CACNA1D, ↑ELAVL2, ↑FABP7, ↓FAT2, ↓L1CAM, ...all 11 |
| LEF1 | transcription regulator | Activated | 2.688 | 7.20E-04 | ↓ACAN, ↑ANXA1, ↑CARD11, ↑CAV1, ↓CD24, ↑CD274, ↑.. ...all 20 |
| IL1B | cytokine | Activated | 2.659 | 5.79E-10 | ↓ACAN, ↓ANGPTL2, ↑ANXA1, ↓APOE, ↑ASS1, ↓BACE1, ...all 80 |
| enzalutamide | chemical drug | Activated | 2.630 | 7.35E-03 | ↑BOK, ↓CYP4V2, ↑ENO2, ↑GADD45G, ↑PPP1R13B, ↑TGF... ...all 8 |
| IL4 | cytokine | Activated | 2.576 | 8.06E-08 | ↓ABLIM1, ↓ACSL6, ↑ADCY1, ↓AGPAT4, ↓ALDH2, ↑ANX... ...all 74 |
| pirinixic acid | chemical toxicant | Activated | 2.547 | 4.69E-03 | ↑APOA2, ↓APOE, ↑BNIP3, ↓CD24, ↑CGA, ↑CTSB, ↓CYP... ...all 24 |
| STAT3 | transcription regulator | Activated | 2.526 | 3.31E-09 | ↓ANGPTL2, ↑BNIP3L, ↑CASC9, ↑CCR7, ↑CD274, ↑CDH5, ...all 58 |
| NR3C2 | ligand-dependent nuclear receptor | Activated | 2.520 | 5.53E-04 | ↑APOC1, ↓APOE, ↓BMP4, ↑CACNA1H, ↑CXCR2, ↑DDIT4, ...all 15 |

| Upstream Regulator | Molecule Type | Predicted Activation State | Activation z-score | p-value of overlap | Target Molecules in Dataset |
|---|---|---|---|---|---|
| ALKBH5 | enzyme | Activated | 2.190 | 1.56E-02 | ↑BNIP3, ↑CAV1, ↑FABP5, ↑FGFR1, ↓FOXO1, ↓IGF2, ↑ND... ...all 9 |
| KLF15 | transcription regulator | Activated | 2.187 | 9.80E-02 | ↑DDIT4, ↑DECR1, ↑FABP5, ↑IRX3, ↑SLC27A1, ↑TNFSF10 ...all 6 |
| methylnitronitrosoguanidine | chemical toxicant | Activated | 2.186 | 3.09E-03 | ↑BTG1, ↑GDF15, ↑KLF10, ↑SERPINE1, ↑TNFSF10 ...all 5 |
| D-glucose | chemical - endogenous mammalian | Activated | 2.176 | 7.05E-05 | ↓ACAN, ↑ANXA1, ↓ANXA3, ↓AQP1, ↑ASS1, ↓BMP4, ↓... ...all 45 |
| IL5 | cytokine | Activated | 2.174 | 9.10E-04 | ↑ASS1, ↑BNIP3, ↑BNIP3L, ↑CCR7, ↓CD24, ↑CISH, ↑CSF... ...all 19 |
| glucocorticoid | chemical drug | Activated | 2.162 | 4.41E-02 | ↑ANXA1, ↑CGA, ↑CYP19A1, ↓GABRP, ↑GBP1, ↓GPR158, ...all 15 |
| ADIPOQ | other | Activated | 2.160 | 7.17E-05 | ↓ANGPTL2, ↑APOA2, ↑APOC1, ↑CGA, ↓CHIC1, ↑CYP19A1 ...all 22 |
| SREBF1 | transcription regulator | Activated | 2.146 | 2.87E-03 | ↑APOA2, ↑BHLHE40, ↑BHLHE41, ↑CTH, ↓CXADR, ↓CYP... ...all 18 |
| IL2 | cytokine | Activated | 2.141 | 5.44E-05 | ↓ACSL3, ↓ACSL6, ↓ADCY3, ↑ANXA1, ↓AUTS2, ↑BHLHE40 ...all 43 |
| TYA-018 | chemical reagent | Activated | 2.138 | 2.57E-03 | ↓ACSL3, ↑BNIP3, ↑CAV1, ↓CHMP6, ↑DECR1, ↑GGH, ↑... ...all 14 |
| IL15 | cytokine | Activated | 2.127 | 1.00E-04 | ↑ANXA1, ↓BAMBI, ↑BTG1, ↑CD274, ↑CD53, ↑CISH, ↑C... ...all 31 |
| cantharidin | chemical drug | Activated | 2.121 | 1.15E-03 | ↓ALDH3A2, ↑BHLHE40, ↑BHLHE41, ↑CYP19A1, ↑ELK3, ↑.. ...all 26 |
| CD38 | enzyme | Activated | 2.111 | 5.35E-04 | ↑ASS1, ↑BNIP3, ↑BNIP3L, ↑CISH, ↑CSF2RB, ↑FAM162A, ...all 14 |
| NEDD9 | other | Activated | 2.111 | 4.95E-05 | ↑BHLHE40, ↑BNIP3, ↑DDIT4, ↑GDF15, ↓ITGA2, ↑PLIN2, ...all 11 |
| IL17A | cytokine | Activated | 2.101 | 1.37E-02 | ↓ACAN, ↑CD274, ↑COTL1, ↓CYP7B1, ↓HMOX1, ↑HSPB8, ...all 23 |
| semaxinib | chemical drug | Activated | 2.093 | 1.20E-07 | ↑AHRR, ↑CARD11, ↑CCR7, ↓CD24, ↑CSF2RB, ↑CYP19A1, ...all 18 |
| lactic acid | chemical - endogenous mammalian | Activated | 2.085 | 2.35E-03 | ↓AGRN, ↑ARRDC3, ↑CACNG4, ↑CD274, ↓CDH3, ↑CPLX1, ...all 16 |
| STAT6 | transcription regulator | Activated | 2.060 | 1.91E-02 | ↓ABLIM1, ↓ALDH2, ↑ANXA1, ↓APOE, ↑ASS1, ↑CCR7, ↓... ...all 24 |
| ethanol | chemical - endogenous mammalian | Activated | 2.056 | 3.83E-01 | ↓BMP4, ↓CYP7B1, ↑ELOVL3, ↑FABP5, ↑FGFR1, ↑GSR, ↑... ...all 18 |
| poly rI:rC-RNA | biologic drug | Activated | 2.035 | 7.78E-04 | ↓ACSL3, ↓AGRN, ↑ANXA1, ↑APOBEC3C, ↓AUTS2, ↑BNI... ...all 44 |
| JAG1 | growth factor | Activated | 2.030 | 5.63E-05 | ↑CYP19A1, ↑GDF15, ↑GPX3, ↑HEY1, ↑HEYL, ↑JAG1, ↑R... ...all 10 |
| 5-fluorouracil | chemical drug | Activated | 2.009 | 1.85E-03 | ↑BNIP3, ↑BNIP3L, ↓CD24, ↑COX6C, ↓EPHA2, ↓FOXO1, ...all 24 |
| CpG ODN 2006 | chemical reagent | Activated | 2.008 | 1.55E-01 | ↑CD274, ↑GBP1, ↑IL2RB, ↑LYN, ↑MVP, ↓MX1, ↑NOTC... ...all 11 |
| 1-palmitoyl-2-(5-oxovaleroyl)-sn-glyce | chemical reagent | Activated | 2.000 | 1.76E-02 | ↓MYOCD, ↑SNAI1, ↑TGFB1, ↑TWIST1 ...all 4 |
| Mrgpra2a (includes others) | G-protein coupled receptor | Activated | 2.000 | 4.00E-02 | ↑CD274, ↑CXCR2, ↑GRINA, ↑TNFRSF1B ...all 4 |
| ITGAV | transmembrane receptor | Activated | 2.000 | 9.85E-02 | ↑AMIGO2, ↑ITGA1, ↑SERPINE1, ↑TGFB1 ...all 4 |
| niraparib | chemical drug | Activated | 2.000 | 1.29E-02 | ↑ECM1, ↑GBP2, ↑RMRP, ↑TNFSF10 ...all 4 |
| lurbinectedin | chemical drug | Activated | 2.000 | 1.51E-03 | ↑JAG1, ↑KLF10, ↑NOTCH2, ↑SAT1 ...all 4 |
| SPEN | transcription regulator | Activated | 2.000 | 1.08E-01 | ↑RPL22L1, ↑RPL30, ↑RPL8, ↑RPS20 ...all 4 |
| MAP2K3 | kinase | Activated | 2.000 | 1.43E-01 | ↑SERPINE1, ↑SGK1, ↑SLC5A5, ↑TNFSF10 ...all 4 |
| M344 | chemical reagent | Activated | 2.000 | 3.21E-02 | ↓BACE1, ↑MAPT, ↑SERPINE1, ↑TGFB1 ...all 4 |

Figure S1: Ingenuity Pathway Analysis (IPA) of Bulk RNAseq data from ARHGAP18 KO compared to WT-Jeg3. A) Graphical representation of predicted interactions. Nearly all observed protein-protein interactions are separated by a grey line indicating "Interaction not predicted". Notable exceptions include RhoA, CDH1, and SMAD4 B) List of upstream regulator predicted to be involved due to expressional changes seen in the dataset. C) List of upstream regulator predicted to be involved due to expressional changes seen in the dataset.
